# CapuchinAI 1.0: Development of a machine learning-based touchscreen paradigm to test cognition in wild capuchins

**DOI:** 10.1101/2025.11.07.687266

**Authors:** Federico Sánchez Vargas, Sai Rakshith Potluri, Jacob Abernethy, Marcela E. Benítez

## Abstract

Advancing the study of primate cognition requires methods that preserve ecological validity while enabling the experimental control typical of laboratory research. We introduce **CapuchinAI**, a field-deployable touchscreen system that integrates real-time facial recognition with automated cognitive testing, providing a novel methodological framework for studying cognition in wild primates. Our approach combines a high-performing (>97% accuracy) YOLOv7-based facial recognition model (*Multiple Capuchins v1.0*) with a portable Raspberry Pi–driven touchscreen–reward apparatus designed for automated operation in natural habitats. The system detects approaching capuchins, initiates video recording, presents touchscreen stimuli, and dispenses food rewards contingent on task performance. During a two-week presentation to two habituated groups of wild white-faced capuchins (*Cebus imitator*) at the Taboga Forest Reserve, **16 individuals** voluntarily interacted with the apparatus, **10 learned to trigger rewards**, and **8 formed and retained robust screen–reward associations**. The rapid habituation and learning rates demonstrate the feasibility of deploying AI-mediated cognitive experiments in the wild. CapuchinAI addresses several long-standing challenges in field cognition research by enabling: (1) autonomous, individualized task administration without researcher intervention; (2) standardized, repeatable trials across individuals and sessions; (3) scalable deployment across groups and sites; and (4) parallel data collection on behavior, identity, and performance. This methodology provides a blueprint for integrating machine learning, touchscreen testing, and automated reward delivery to study within- and between-individual cognitive variation under natural conditions. CapuchinAI represents a significant step toward long-term comparative research on primate cognition by making laboratory experimental paradigms accessible in the wild, bridging the gap between lab and field.

**Research Highlights:**

- We present CapuchinAI, a field-ready touchscreen testing station that uses real-time facial recognition to study cognition in wild capuchin monkeys
- We developed a YOLOv7-based facial recognition model (Multiple Capuchins v1.0) that identifies individual capuchins with >97% precision and recall from static images, video, and live footage, enabling fully automated, individualized testing in the wild.
- We integrated a version of this model into a closed-loop touchscreen–reward pipeline that detects an approaching monkey, presents a basic learning task, and automatically delivers food rewards based on the monkey’s responses.
- Wild capuchins rapidly habituated and learned touchscreen–reward associations, showing that AI-enabled touchscreens provide a scalable field method for deploying lab-style cognitive tests and mapping individual differences across tasks, species, and sites.

## 1. INTRODUCTION

Primates stand out from other mammals by having unusually large brains and advanced cognition (Roth & Dicke, 2012). This has established primates as key models for the study of cognitive evolution, with their complex and diverse social world, extended development and life history, and species-specific ecological adaptations marking them as prime candidates to study how brains and minds evolve and adapt (Dunbar & Shultz, 2007; Grabowski et al., 2023; Leigh, 2004). Understanding primate cognitive evolution is an interdisciplinary objective, uniting anthropologists focused on the pressures that have selected for larger brains with psychologists exploring the mechanisms driving cognitive choices. A central assumption in both approaches is that cognitive abilities are adaptive (Byrne, 2000; Dukas, 2004): individuals who are more effective at acquiring, processing, and using information to make informed decisions should be more likely to survive and reproduce (i.e., fitness). Yet, despite this foundational premise, surprisingly few studies have directly linked cognitive performance to real-world fitness outcomes in primates (Huebner et al., 2018). One primary reason for this gap is methodological. Cognition has traditionally been studied in captivity, where experimental control and replicability allow researchers to isolate the mechanisms underlying complex behaviors (Shettleworth, 2001). While these settings provide experimental precision, they are necessarily detached from the rich social and ecological contexts in which primate cognition evolved. Consequently, it is difficult to distinguish what primates *can do* in the laboratory from what they *actually do* in nature. Field studies, by contrast, capture the ecological validity and functional significance of cognitive traits but often lack the experimental rigor and control necessary to infer underlying mechanisms. The strengths of one approach, therefore, highlight the weaknesses of the other: captive experiments provide control but lack ecological validity, field studies provide ecological validity but lack controls.

Field experiments have begun to narrow the gap between captive and wild research by introducing shared frameworks for testing cognition under natural conditions (reviewed in Benítez et al., 2022). Approaches such as artificial-fruit “puzzle boxes” and field-adapted foraging tasks have provided valuable insights into how primates innovate, learn, and cooperate in ecologically relevant contexts (e.g., van de Waal et al., 2010; Canteloup & van de Waal., 2020; Amici et al., 2020; Molesti & Majolo 2016). These studies demonstrate that interactive experiments are feasible in the wild and can yield meaningful data on social learning, problem-solving, and cooperation. Yet they remain constrained in scale and resolution: participation is often unpredictable, dominant individuals can monopolize access, and most designs allow only single interactions, limiting replication, sample size, and standardization (Lopresti-Goodman et al., 2022; Thornton & Samson, 2012). As a result, while these field experiments have been pivotal in extending cognitive research beyond captivity, they have yet to yield the long-term, fine-grained data needed to quantify individual variation across multiple cognitive domains —a critical step in understanding the evolution and adaptive significance of primate cognition.

Here, we describe a novel method to study cognition in the wild. Our “smart testing stations” integrate advances in machine learning with a field-friendly touchscreen interface to investigate cognition in wild white-faced capuchins (*Cebus imitator*) at the Taboga Forest Reserve in Costa Rica. Recent advances in machine learning and computer vision now make it possible to automatically recognize and track individual primates in natural environments (Schofield et al., 2019). These tools have already been used to reconstruct social networks and quantify the stability of individual social tendencies in wild chimpanzees (*Pan troglodytes*; Schofield et al., 2023), offering unprecedented opportunities to monitor behavior at scale. Building on these advances, our rugged testing stations combine machine-learning–based identification with a remotely operated touchscreen apparatus that can tailor cognitive testing to each individual in real time. This design is equipped with a 3D-printed food dispenser that dispenses 32 rewards (in our case, dried forest bananas), enabling controlled, repeatable trials within natural group settings. This integration overcomes key logistical and methodological barriers in field cognition research—allowing for the automated, high-resolution collection of multi-timepoint data on individual performance across multiple cognitive domains (Benítez et al., 2022).

We see several immediate and long-term goals for this design. In the short term, it enables researchers to (1) assess individual variation across multiple domains of cognition, (2) test whether performance in one domain predicts performance in others, and (3) examine how such variation relates to how successfully individuals navigate social and ecological challenges. Over the long term, our goal is to (1) systematically track within-individual changes in cognitive performance across life history stages, and (2) generate one of the most comprehensive longitudinal datasets on cognition in a wild primate population. By pairing long-term cognitive data with long-term demographic, behavioral, ecological, and hormonal data, we can begin to connect individual variation in cognitive trails to fitness-relevant outcomes such as social success, reproduction, and survival. This methodology fills a critical gap, offering a scalable, field-ready approach for systematically testing multiple domains of cognition in the wild and linking cognitive variation to the evolutionary processes that shape it.

In this paper, we detail the methodology of this new approach, including the facial recognition model, data processing, model pipelines, and touchscreen design. Specifically, we address three key questions: (1) Can we develop a high-quality facial recognition model for capuchins? (2) Can we create a field-friendly touchscreen device that runs this model in real time to deliver and adapt cognitive experiments, controlling task presentation and reward? And (3) will wild capuchins reliably interact with this system? Together, these steps outline a practical roadmap for integrating artificial intelligence and experimental cognition in the wild, providing, for the first time, a systematic way to evaluate and monitor the cognitive abilities of wild primates.

### 1.1 CapuchinAI: A new paradigm for studying wild capuchin cognition

Capuchin monkeys are a key species for studying comparative cognition. They share several traits with humans, such as large brains relative to body size (Rilling and Insel, 1999), extensive cooperation both in captivity (Brosnan, 2011; Brosnan et al., 2006) and in the wild (Perry, 1996; Scarry, 2017), prosocial behaviors (Brosnan, 2011; Claidière et al., 2015), and sophisticated skills in extractive foraging (Barret et al., 2018). Since capuchins and humans diverged approximately 23 million years ago (Glazko & Nei, 2003), they serve as a valuable comparative model to explore the social and ecological factors that selected for the evolution of larger brains. Compared to many other non-human primates, both tufted capuchins (*Sapajus apella* spp.) and white-faced capuchins (*Cebus capucinus* spp.) are particularly skilled at innovation, learning, and solving complex problems. In captivity, tufted capuchins are easily trained on a range of manual and computer-based tasks (e.g., Parrish et al., 2015), providing a detailed understanding of their cognitive capabilities. However, because most of what we know about capuchin cognition comes from captive studies mainly of *Sapajus*, there is still much to learn about how they use their large brains in natural environments.

White-faced capuchins have been studied extensively in the field, yielding rich descriptions of their social structure, cooperation, extractive foraging, and social learning capabilities. What remains comparatively unclear are the cognitive mechanisms underlying these behaviors: how individuals acquire, represent, and use information to make decisions, and how such variation maps onto socioecological conditions. One reason is practical. Unlike their *Sapajus* relatives, white-faced capuchins are not readily studied in captivity, which has constrained experimental work and left a gap between documented behavioral observations in the wild and direct measures of cognition.

The white-faced capuchins at the Capuchinos de Taboga field station in the Guanacaste region of Costa Rica represent an ideal population to begin such an investigation. This site contains the highest-density population of white-faced capuchins recorded in the wild (Tinsley Johnson et al., 2020). At this site, four groups have been fully habituated and are subject to daily behavioral observations and hormone sample collection. Furthermore, three of these groups have also been habituated to testing platforms installed in their home ranges, where they voluntarily participate in multiple non-invasive cognitive experiments. These testing platforms have been used to conduct several cognitive experiments, generating a large, multi-year dataset of high-resolution videos and images of individuals who frequent the platforms. These videos provided the dataset for training our models (**Section 2.1**), and these platforms provide the foundation for deploying smart testing stations (**Section 3.1**).

Our main objective is to develop and implement “smart” testing stations for long-term cognitive assessment of the wild capuchins at Taboga. Each station uses a machine learning facial-recognition model to detect an approaching monkey, identify the individual in real time, and retrieve the “bookmark” for where that individual left off in the cognitive testing battery. The station then presents the appropriate next task and level on the touchscreen interface, automatically guiding subjects through a scaffolded cognitive battery of tasks (see **Figure 1** for a visual representation of this workflow).

**Figure 1.**
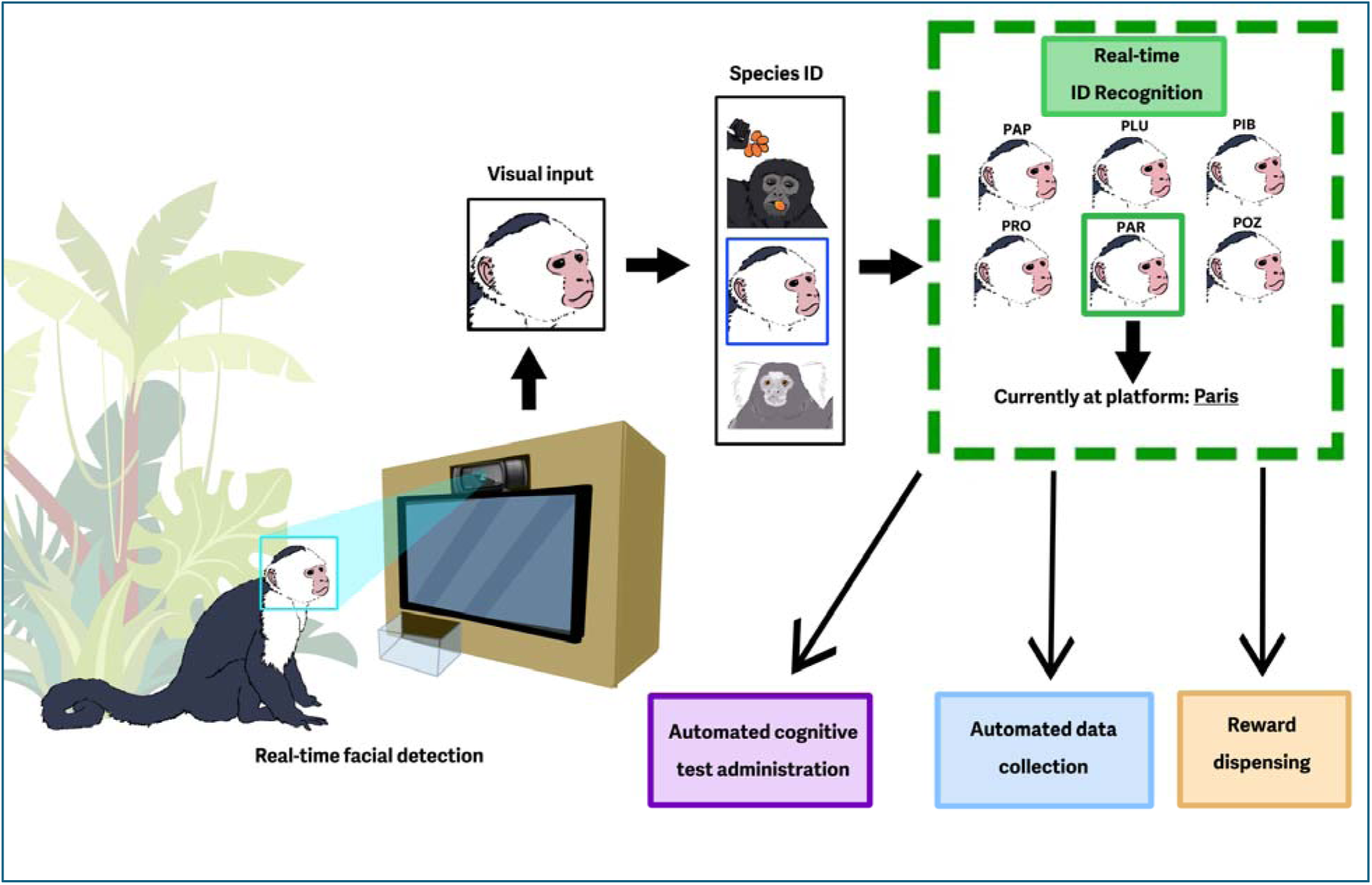
Diagram of real-time, automated facial detection and cognitive testing touchscreen apparatus: As a monkey approaches the apparatus (left), the machine learning facial recognition model detects the presence of a potential participant. The model determines that the approaching animal is a capuchin and then, in real time, recognizes the individual’s identity based on its training dataset. With this information, the model can then automatically administer touchscreen cognitive tasks, collect and store data, record video of the participating monkey, and administer food rewards for correct responses. Note that, since the model runs inferences at more than 30 frames per second, any sudden change in the identity of the participating individual will seamlessly trigger downstream changes in the stimuli presented and the data collected. Illustration based on testing conditions at Taboga Forest Reserve, Costa Rica.

This design offers two central advantages. First, our AI facial recognition software enables autonomous, individualized cognitive testing, allowing researchers to manage participation remotely and adjust task difficulty to each monkey’s performance, so each individual progresses at their own pace. In typical lab workflows, researchers manually program a station for a single, preselected subject, which is not feasible in the wild. Our approach preassigns tests for all potential participants before a session begins. As individuals arrive and depart, the integrated facial-recognition system detects both the presence of a capuchin and its specific identity in real time, then serves the individualized task list to the animal at the screen. Because detection and identification refresh continuously at millisecond resolution, changes in participants—for example, when one individual displaces another—do not disrupt testing; the software immediately switches to the tests preassigned to the new participant.

Second, the touchscreen interface will permit us to administer cognitive tests already employed and highly validated in lab settings (e.g., Richardson et al., 1990; Butler et al., 2019) and, more recently, in wild vervet monkeys (Harrison et al., 2023). These include simple association formation paradigms (**Section 3**), delayed match-to-sample tasks to test working and long-term memory (Colares Leal et al., 2020; Sosnowski & Brosnan, 2025), ‘go/no-go’ tasks to test impulse control (Picanço & Barros, 2015), and rule reversal tasks to test cognitive flexibility, among others.

Between July and August 2025, we piloted the initial version of our machine learning-based touchscreen cognitive testing device, ‘**CapuchinAI v1.0,’** with two groups at Taboga Forest Reserve. Below, we explain the development and operation of the software (**Section 2.1**) and hardware (**Section 2.2**), and demonstrate how wild capuchin monkeys habituated and interacted with the device (**Sections 2.3, 3.1 – 3.2**). We conclude by outlining future development steps (**Section 3.3**) and discussing the strengths, limitations, and potential for adaptation to other primate species and field stations (**Section 4**).

## 2. DESCRIPTION

Over the past decade, the computer vision community has invested much effort into a new class of models, known as *deep convolutional neural networks* (CNNs), one of the staples of deep learning (LeCun et al., 2015). There has been significant progress in the success of ML-based object detection in recent years, with the best models achieving essentially human-level performance on computer vision tasks (Wang & Deng, 2021). This has been driven in large part by methodological improvements, aided by massive advances in chips designed for learning (i.e., GPU technology) and the vast amount of available data.

Recently, there has been growing interest in using deep learning models to detect animal species and to deploy them in conjunction with camera traps in the wild (Norouzzadeh et al., 2018; Tabak et al., 2018). In primates, machine learning has been used to successfully identify species and individuals within a species, for example, red-bellied lemurs (*Eulemur rubriventer,* Crouse et al., 2017), rhesus macaques (*Macaca mulatta,* Witham, 2018; Zhang et al., 2025), and golden snub-nosed monkeys (*Rhinopithecus roxellana,* Guo et al., 2020). However, nearly all this work is based on data collected in highly structured environments, often with monkeys in captivity, or with stationary camera setups. A key exception is fully automated, face-based identification of wild chimpanzees from long-term field video (Schofield et al., 2019), which demonstrates the feasibility of AI models under field conditions. Building on this work, our model adds two capabilities to the current AI facial recognition work in primates: 1) a reliable operation in arboreal, forest habitats with shifting light, occlusion, and rapid movement, and 2) millisecond-level identity resolution, allowing the model to recognize multiple individuals in real time.

### 2.1. Can we develop a high-quality capuchin facial recognition model?

The first step was to create the dataset for training a precise, accurate, and robust capuchin facial recognition model. At Capuchinos de Taboga, we gathered an extensive collection of high-quality close-up GoPro videos of capuchins interacting with cognitive experiments on testing platforms. Each video was manually coded, identifying capuchins in each frame and their behaviors of interest using an open-source event-logging software, Behavioral Observation Research Interactive Software (BORIS; Friard & Gamba, 2016). Using the timestamped BORIS output, we configured a Python script to extract high-quality frames from these videos every second that a capuchin was present. Once frames were extracted, they were stored as individually labeled images.

These images were then uploaded to Roboflow (Dwyer et al., 2025), where we manually placed bounding boxes around each capuchin’s face to define the region of interest. Roboflow is an advanced tool designed to streamline and optimize the process of preparing datasets for machine learning, particularly in the field of computer vision. To ensure our model was trained on a balanced dataset, we applied an additional sampling step after frame extraction. This post–frame extraction sampling ensured that each subject contributed an equal number of frames to the final dataset, preventing any single subject from dominating the training data. To bolster model robustness and ensure strong performance under varying lighting and color conditions, we used Roboflow’s dataset augmentation tools to generate altered versions of the training images. These augmentations modified lighting, resolution, and image blur to create additional examples for the model that mimic the variability of filming conditions in the wild. The final dataset consisted of 2,160 unique images — over 4000 images with the augmented dataset — evenly distributed across six adult male capuchins (Pery, Pirulo, Pele, Pesto, Pio, and Pedro).

For our model, Multiple Capuchins v1.0, we used Ultralytics YOLOv7 (You Only Look Once, version 7-nano) object detection model architecture (Wang et al., 2023). This architecture is specialized for rapid, real-time object detection, making it well-suited for high-accuracy identification and classification of objects in live video streams. The model was trained with a batch size of 8 for 300 epochs. All settings were defined in a YAML file (e.g., class names, number of classes, and paths to the training and validation sets). The Multiple Capuchins v1.0 model performed exceptionally well on the held-out test dataset (images the model was not trained on), including perturbed images, achieving 97.5% precision and 98.2% recall. In practical terms, this means that when the model was presented with a capuchin face—even when lighting or resolution were altered—it detected and attempted to classify that face 98.2% of the time, and when it did so, it assigned the correct monkey ID 97.5% of the time. We found that Multiple Capuchins v1.0 can successfully classify static images and pre-recorded videos (**Figure 2**). The next step was to figure out how to integrate a version of this model to run on a low-energy, field-friendly computer interface.

**Figure 2.**
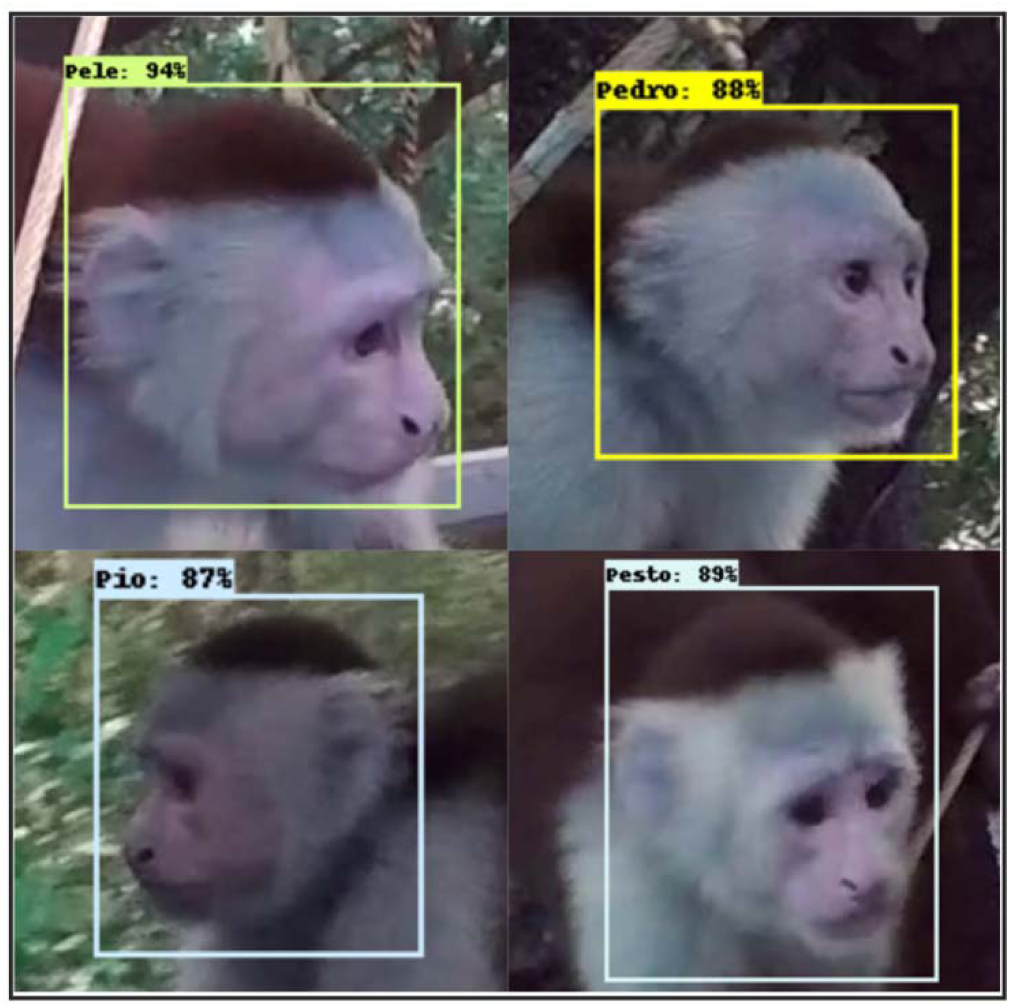
MultipleCapuchins v1.0 Facial Recognition Model Performance: Sample still images taken from output videos of the MultipleCapuchins v1.0 machine learning model performing capuchin facial detection. This model was developed on Ultralytics YOLOv7 architecture and trained on the identities of six male capuchins from video footage collected during previous cognitive experiments at Taboga Forest Reserve, Costa Rica. Bounded boxes indicate the area where the model recognizes a capuchin face. Names and percentages in the upper left corner of the bounded boxes indicate the model’s guess and confidence in the identity of the individual within the box. Note that the four individuals pictured here are each looking in different directions under distinct lighting conditions – yet the model retains high confidence.

### 2.2. Can we create a field-friendly touchscreen device that runs this model in real time to deliver and adapt cognitive experiments, controlling task presentation and rewards?

Having developed a highly accurate capuchin facial-recognition model, our next step was to build a field-friendly touchscreen apparatus that could run live facial detection, present tasks to interacting monkeys, and dispense food rewards. Our apparatus, **CapuchinAI v1.0**, runs on a Raspberry Pi 5 (RPi5), a compact Linux-based computer well suited for multi-input/output applications that must operate reliably in remote locations (e.g., primatology field sites) (Mathe et al., 2024).

CapuchinAI, controlled by the RPi5, integrates the following components:

1. A 15-inch screen to display stimuli to participating monkeys.
2. An infrared touch overlay that converts the screen into a touch display. This type of touchscreen is ideal for work with primates because it is inexpensive, durable, and can be activated with any part of the primate’s body (e.g., finger, whole hand, mouth, nail, tongue).
3. A high-definition USB webcam that provides a live video feed for the model to run detection on.
4. An electronic relay device, connected to a 3D-printed food reward dispenser, that controls reward delivery by closing a circuit for a defined period whenever a monkey touches the screen.

The entire setup is powered by two low-cost, widely available travel power banks (one for the display and one for the RPi5) and a 12-volt battery pack for the food dispenser’s motor. Initial tests showed that the apparatus could run uninterrupted on this power supply for at least 8 hours. It is encased in a wooden frame measuring (19 × 21× 5 inches) that fits into the cognitive testing platforms at Taboga and is internally organized to allow easy access to all components, facilitating on-the-fly adjustments (e.g., adjusting screen or camera placement, replacing power sources, refilling the food dispenser). **Figure 3** provides a visual schematic of this setup.

**Figure 3.**
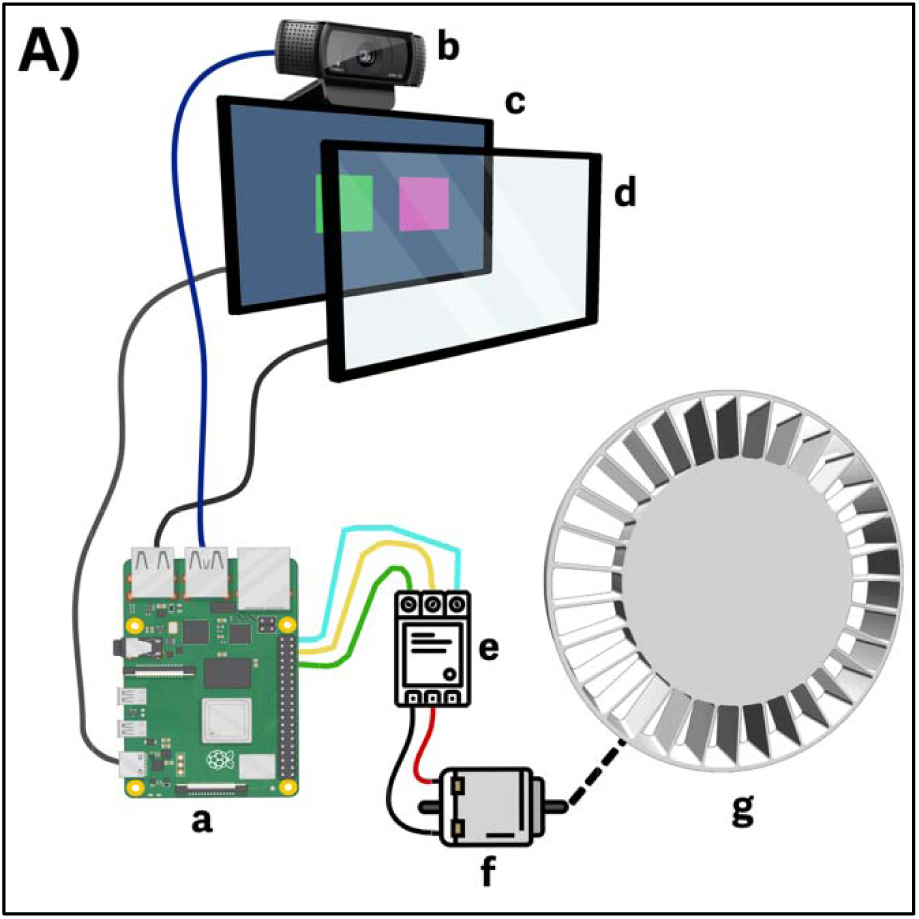
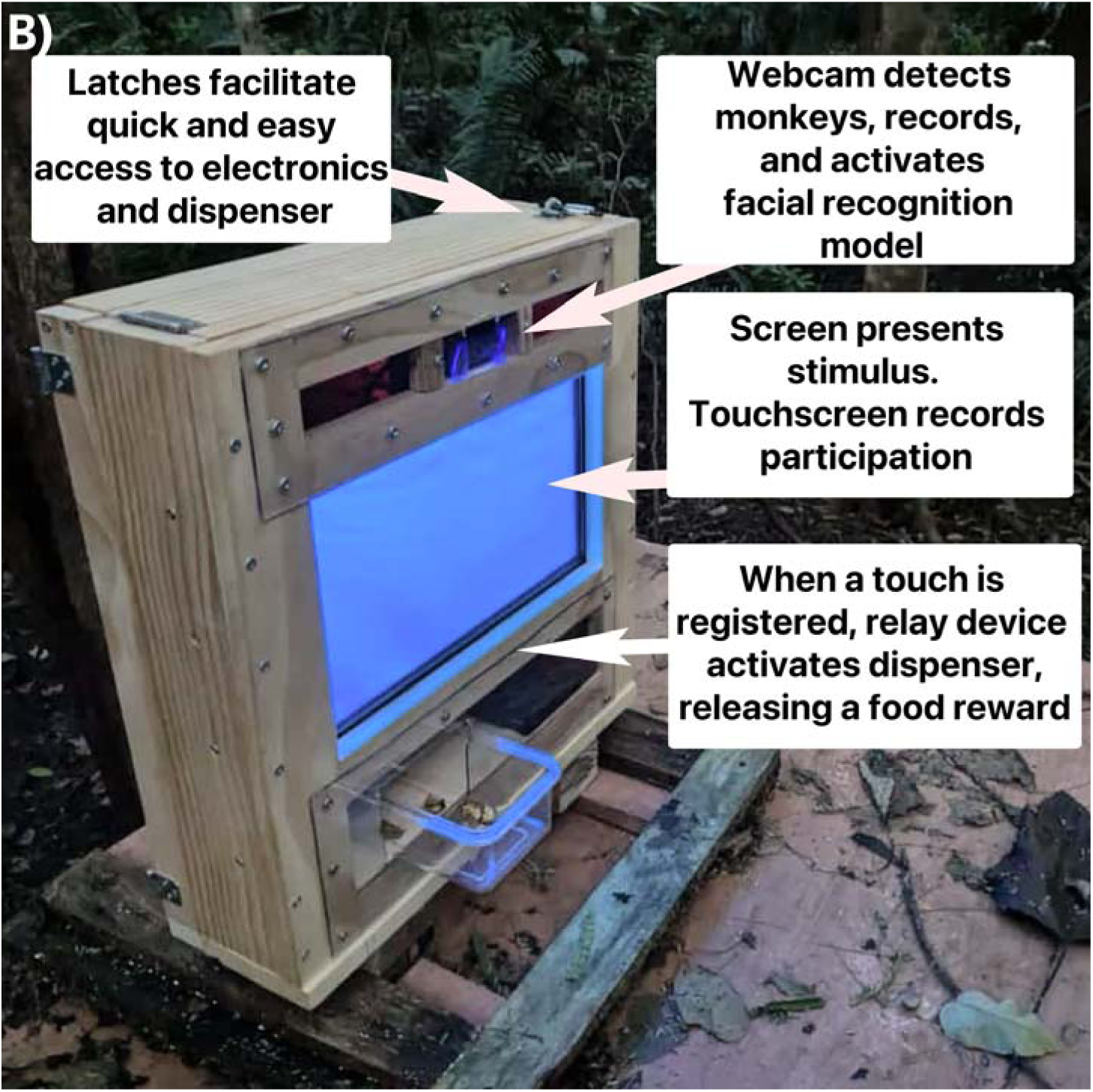
CapuchinAI v.1.0 Cognitive Testing Apparatus. **A) Diagram of functional components of apparatus:** CapuchinAI v.1.0 integrates multiple peripheral components through a series of python scripts run on a Raspberry Pi 5 (**a**). The Raspberry Pi 5 connects via USB to a full-HD webcam (**b**), which feeds live video footage to a continuously running facial recognition machine learning model. Simultaneously, cognitive testing visual stimuli are presented to primate participants via a display screen (**c**), and participants’ screen touches are recorded via an infrared touchscreen overlay (**d**). If a participant successfully completes an assigned task, the Raspberry Pi 5 sends a signal to a relay device (**e**) via its GPIO (general purpose input/output) pins. This relay device completes an electrical circuit powering a 12-volt DC motor (**f**) for a defined amount of time, allowing a rotary food reward dispenser (**g**) to rotate exactly the degrees needed to release one food reward. Note: components not drawn to scale. **B) CapuchinAI v.1.0. apparatus deployed at testing platforms in Taboga Field Site, Costa Rica**. The image shows the completed apparatus ready for interaction by primate participants. Key visible components are described.

For our prototype, we deployed a modified version of our Multiple Capuchins v1.0 model on the RPi5, reimplemented using the Ultralytics YOLOv5 architecture. YOLOv5 is a lighter weight, widely used model that has been successfully run on Raspberry Pi platforms, making it a practical choice for efficient, real-time inference in our field setting. In this prototype, we simplified the model detection categories to detect any capuchin face rather than classify individual identities, as the model would be encountering capuchins it was not trained on. This design served two goals: first, it enabled the system to trigger video recording whenever any capuchin approached the touchscreen, rapidly generating a larger, more up-to-date dataset of faces from interacting monkeys; second, it allowed us to offer the touchscreen task to all habituated capuchins, not only those included in the original YOLOv7 training set. Because the identities of all participating individuals are already known from long-term monitoring, the resulting videos and images can be used to train the next, more comprehensive version of our individual-level facial recognition model (see **Section 3.2**).

To integrate our facial recognition model with our apparatus, we developed two Python scripts. The first of these, “**capuchin_recorder.py**”, initializes the webcam and starts running detection on the camera’s live feed at approximately 30 frames per second, providing high-resolution, real-time facial detection. Since our goal at this stage was to collect videos of all individuals approaching the platform, this script instructs the camera to start recording when face detection occurs and to stop recording after 10 seconds elapse with no capuchins in view of the camera. This loop continues for as long as the script runs, generating multiple videos per session without quickly overloading the RPi5’s storage or processing capacity. At the end of the session, the script places all recorded videos in a designated folder, along with a log of the start and end times for each video.

The second Python script, “**touchscreen_reward_interface.py**” uses pygame v.2.6.0 (McGugan, 2007), a python library designed to facilitate the creation of interactive stimuli for use in games, cognitive testing, or other user-facing interfaces. Using pygame, we developed a simple habituation stimulus for monkeys: a screen-wide blue square that records when an interacting monkey touches it. When a touch event occurs, our script signals the relay device to close the dispenser’s motor circuit, triggering the automatic dispensation of a food reward. The screen next flashes black for 2 seconds during a cooldown period, then returns to blue, signaling that it is ready to register a new touch.

These two Python scripts are integrated to run simultaneously and share key information. Critically, the stimulus presentation and reward dispensing script only signals the relay device to deliver a reward if a capuchin face has been detected. As a result, the system prevents rewards from being dispensed to non-study species (e.g., other species that may interact with the apparatus) and provides the foundation for individual-level control over the number of trials each monkey can complete per day, a promising aspect of our methodology that we elaborate in **Section 4**.

## 3. EXAMPLE

### 3.1. Will wild capuchins reliably interact with this system?

Between July and August 2025, we presented our CapuchinAI v1.0 prototype to two groups of wild white-faced capuchins at the Taboga Forest Reserve that have been under near-daily observation by the Capuchinos de Taboga project since 2017. These two groups of capuchins, Tenori (N = 25 individuals) and Palmas (N = 26 individuals), are fully habituated to testing platforms within their home ranges and have participated in ongoing cognitive tasks. The apparatus was secured on testing platforms opportunistically, depending on the focal group’s location on any given day. The setup was always conducted outside the view of any capuchin groups to prevent associations between the researchers and the food rewards. Food rewards consisted of sliced, dried banana pieces, a natural food at Taboga that these capuchins readily eat. At this stage, our goals were to habituate monkeys to interact with the new apparatus voluntarily and to assess the performance of the real-time capuchin face-recognition model in the field, both by triggering video recordings to generate training data for an expanded model and by driving the automated stimulus–reward pipeline described above.

During our brief pilot habituation phase, we conducted six sessions with each group. Throughout all sessions, 16 monkeys (see Table 1 & Table 2) interacted with the testing apparatus, while an additional 14 individuals observed these interactions. All individuals who engaged with the apparatus displayed exploratory behaviors such as touching, licking, sniffing, and/or aggressing the touchscreen station. Of the 16 capuchins that interacted with the testing stations, 10 learned to trigger the reward mechanism by touching the screen, and each of these continued to make exploratory touches that yielded additional rewards. Eight capuchins demonstrated behaviors consistent with having formed a full association between touching the screen and receiving a reward (5 in Palmas group and 3 in Tenori group), with one additional individual from Tenori possibly having formed this association (though we could not confirm as rewards had run out). We defined a full association as being established when an individual 1) directed most or all touches towards the screen area, and 2) showed anticipatory behaviors, such as turning to face the dispenser opening before or during touching the screen, or reaching for it immediately after a deliberate touch, prior to receiving a reward. For five of these individuals, we also observed that the association was retained after a week of no interaction with the apparatus. A brief narrative description of each individual’s behavior across sessions for Palmas group can be found in Table 1, and for Tenori group in Table 2.

**Table 1.** Participants from ‘Palmas’ Group in Habituation Sessions for Touchscreen Cognitive Testing: The first column lists the names of individuals who interacted directly with our testing apparatus during at least one of six sessions. The second column contains the sex and age class of each participating individual. The remaining columns contain a description of the behavior of each individual across sessions. If an individual did not arrive at testing platforms on a given session, they are marked as **N/A**. **Exploratory behavior**: behavior in which individuals approached and investigated the apparatus, without directing special attention to the screen. **Exploratory touches**: deliberate touching and interaction with the screen, without indication of a complete association of screen touching with reward dispensation. **Full association formed**: individual directs touches fully or predominantly at screen and anticipates reward dispensing after touching screen. **Full association retained**: an individual who has previously formed an association maintains that behavior immediately upon returning for a new session.

| Individual | Sex/Age class | Session 1 | Session 2 | Session 3 | Session 4 | Session 5 | Session 6 |
| --- | --- | --- | --- | --- | --- | --- | --- |
| PAPI | Male, | Exploratory behavior, no | N/A | Full association | Full association | N/A | Full association |

*Table 1 — Participants from ‘Palmas’ Group in Habituation Sessions for Touchscreen Cognitive Testing:* The first column lists the names of individuals who interacted directly with our testing apparatus during at least one of six sessions.
|  | Alpha | screen touch |  | **formed** | **retained** |  | **retained** |
| --- | --- | --- | --- | --- | --- | --- | --- |
| PLUTO | Male, Adult | N/A | N/A | N/A | N/A | N/A | Exploratory touches, observes PIBE touching screen and steals food rewards, **full association formed** |
| PIBE | Male, Subadult | N/A | N/A | Exploratory touches, close observation of PAPI | **Full association formed** | N/A | **Full association retained** |
| PARIS | Male, Subadult | N/A | N/A | N/A | N/A | Exploratory behavior, no screen touches | Exploratory touches, **Full association formed** |
| PROBOSCIS | Male, Juv. | N/A | Exploratory behavior and screen touch. | Exploratory screen touches, displaced by PAPI and PIBE | Exploratory screen touches, observing PAPI and PIBE | **Full association formed** | N/A |
| POZA | Male, Juv. | N/A | N/A | Exploratory behavior, no screen touches | N/A | N/A | N/A |

**Table 2.** Participants from ‘Tenori’ Group in Habituation Sessions for Touchscreen Cognitive Testing: The first column lists the names of individuals who interacted directly with our testing apparatus during at least one of six sessions. The second column contains the sex and age class of each participating individual. The remaining columns contain a description of the behavior of each individual across sessions. If an individual did not arrive at testing platforms on a given session, they are marked as **N/A. Exploratory behavior**: behavior in which individuals approached and investigated the apparatus, without directing special attention to the screen. **Exploratory touches**: deliberate touching and interaction with the screen, without indication of a complete association of screen touching with reward dispensation. **Full association formed**: individual directs touches fully or predominantly at screen, and anticipates reward dispensing after touching screen. **Full association retained**: individual that has previously formed association maintains that behavior immediately upon returning for a new session.

| Individual | Sex/Age class | Session 1 | Session 2 | Session 3 | Session 4 | Session 5 | Session 6 |
| --- | --- | --- | --- | --- | --- | --- | --- |
| PEDRO | Male, Alpha | N/A | N/A | Exploratory behavior, accidental screen touches | Accidental screen touches, steals food rewards from TSUNAMI | Exploratory touches on screen. | N/A |
| TROMPUDO | Male, Adult | Multiple exploratory touches to screen | Exploratory touches on screen | **Full association formed** | N/A | **Full association retained** | **Full association retained** |
| TEMPISQUE | Male, Subadult | Exploratory Behavior | N/A | **Full association formed** | **Full association retained** | **Full association retained** | **Full association retained** |
| TSUNAMI | Male, Subadult | Exploratory behavior | N/A | Exploratory touches, deliberate | **Full association formed** | N/A | **Full association retained** |
| TILARAN | Male, Subadult | Multiple exploratory touches to screen | N/A | Exploratory touches, deliberate | No touches on screen | Exploratory touches on screen | N/A |
| TACO | Male, Subadult | N/A | N/A | Exploratory behavior | Exploratory touches on screen | N/A | Purposeful touches, but dispenser ran out of rewards. |
| TOBIAS | Male, Juv. | Exploratory behavior | N/A | N/A | N/A | N/A | N/A |
| TIQUICIA | Male, Juv. | Exploratory behavior, rapidly displaced | N/A | Exploratory behavior, attempts touching screen but is displaced | N/A | N/A | N/A |
| TAMARINDO | Female, Adult | N/A | N/A | Arrived at platform and approached apparatus, but was displaced before she could interact. | N/A | N/A | N/A |
| TURRIALBA | Female, Juv. | Exploratory behavior, rapidly displaced | N/A | N/A | N/A | N/A | N/A |

Taken together, the rates of voluntary participation and rapid habituation of multiple individuals across both groups in the short timeframe in which they encountered our experimental apparatus (2 weeks per group) suggest that this is a highly viable method to study cognition in the wild in our model system — and easily adaptable to other primate species and field sites. Furthermore, participating individuals rapidly learned an association between touching the screen and receiving a food reward. They were further highly motivated to participate whenever they were near the testing apparatus, suggesting that this method may indeed be used to bring true-and-tested lab paradigms to the field. The motivation of individuals to continue participating voluntarily across sessions is promising and suggests that with more prolonged exposure to the apparatus, we’ll be able to assemble a dataset comparable to captive cognition studies.

### 3.2. Ongoing development of CapuchinAI v2.0 — Individual facial recognition and personalized cognitive testing

The deployment of the CapuchinAI v.1.0 touchscreen apparatus succeeded not only in habituating multiple individuals to using the touchscreen to earn a food reward, but also employed our simplified Multiple Capuchins v.1 facial recognition model to generate an abundance of video clips of monkeys interacting with the apparatus (**Figure 4**). Our facial recognition script worked as intended, initiating recording via the connected camera when a detection event occurred and stopping after 10 seconds without any faces detected. This resulted in a library of footage of all 16 participants across both tested groups.

**Figure 4.**
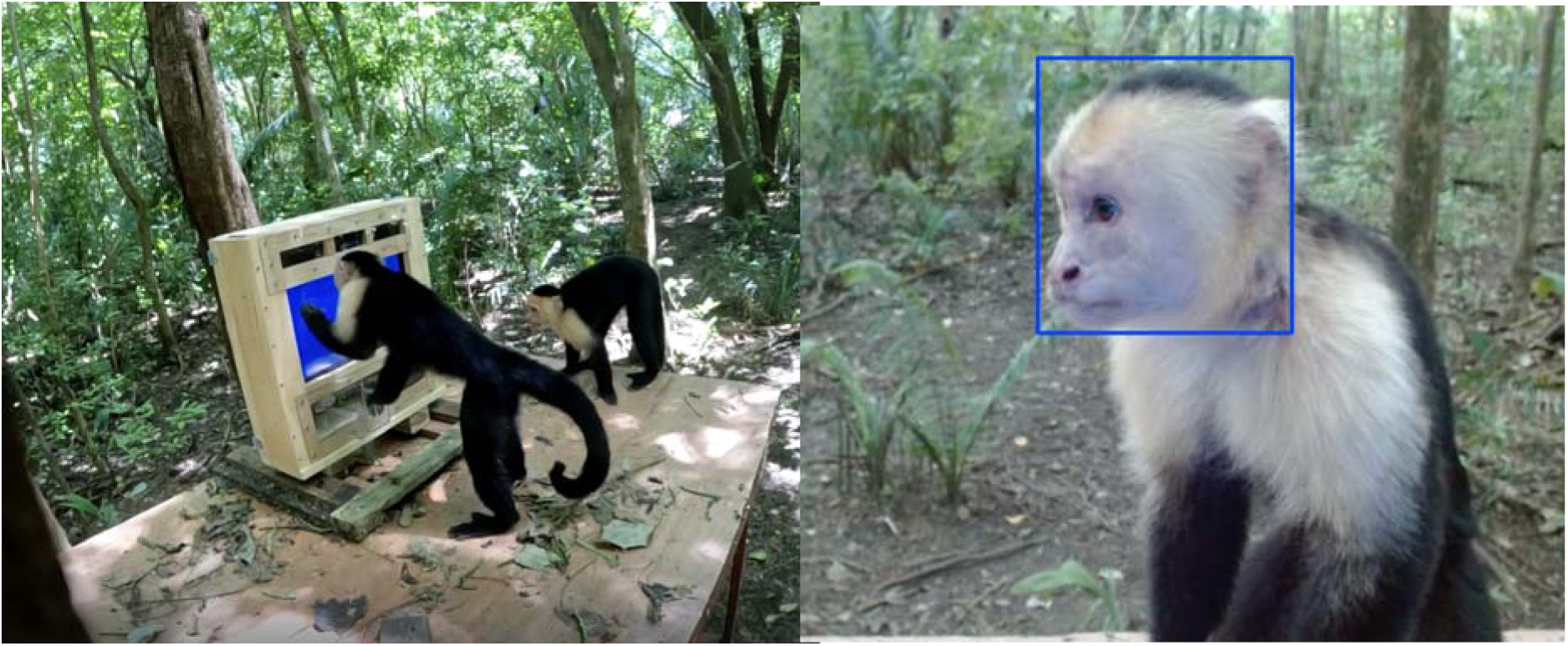
Habituation to testing apparatus and video collection via facial recognition model. **Left:** PAPI (alpha male in Palmas group) purposefully touches the screen with his left hand while placing his right hand on the food dispensing area and looking down at it, displaying an association between touch and reward dispensing. PIBE (subadult male) observes closely. **Right**: still image of PAPI taken from video recorded by CapuchinAI v.1.0’s HD webcam. Blue bounding box indicates that the facial recognition model is actively detecting a capuchin face within the designated area. Extracted frames such as this will be used to train the next iteration of our facial recognition model.

To reduce the effort involved in manually extracting frames from these videos, we will use the same facial recognition model to scan each frame and automatically extract all frames that contain a face through a third Python script. These frames are currently being labeled to train the next version of our facial recognition system, Multiple Capuchins v.2.0. While the initial model achieved high accuracy in detecting and labeling six monkeys, the new model will be able to distinguish all 16 monkeys that participated in our pilot deployment. This updated facial recognition model will allow us to achieve our goal of acquiring individual-specific cognitive data over time across multiple domains of cognition using touchscreen testing methods validated in captive settings (e.g. Colares Leal et al., 2020; Fagot & Cook, 2006; Picanço & Barros, 2015). Our efforts so far have demonstrated that it is possible to **a**) develop a high-quality AI model for individual primate facial recognition, **b**) implement this model for real-time detection in wild settings, and **c**) use the model to regulate the administration of stimuli and dispensing of rewards for voluntarily participating wild primates.

In the next version of our pipeline, we plan to implement a feature that uses the updated facial recognition model to both provide individualized testing <u>and</u> record videos of approachi*ng* individuals. When the model confidently recognizes a capuchin it was trained on, it assigns a specific cognitive test. If the model’s confidence is low—meaning the individual is unfamiliar—it assigns the baseline habituation stimulus and starts recording video. These recordings will be used to periodically update the model, helping it recognize new monkeys as they interact with the testing setup.

## 4. COMPARISON AND CRITIQUE

### 4.1 CapuchinAI provides an unprecedented level of experimental control in the wild

There is a classic trade-off in studies of primate cognition between ecological validity and experimental control. Studies in the wild capture behavior in the fitness-relevant contexts in which cognition evolves, but it has historically been difficult to address the mechanisms underlying observed patterns, something that laboratory experiments do well. The method we describe here takes sizeable steps toward bringing the best of the lab to the field, increasing our control over cognitive data collection in several ways.

One major difficulty in studying cognition in the wild is that experiments have typically been limited to one trial per session (e.g., Lopresti-Goodman et al., 2022; Thornton and Samson, 2012), which restricts the number of individuals that can be tested at one time and increases the likelihood that a single individual will monopolize access (Benítez et al., 2022). Our method directly addresses this limitation. By using machine vision, the apparatus can rapidly shift tests from one identified individual to another and allow multiple individuals to participate in a single session. Because the system does not rely on an experimenter to switch tasks based on participant identity, but instead automates recognition of the participant, assignment of the appropriate experiment, and reward delivery, multiple apparatuses can be installed in close proximity without any one individual monopolizing testing opportunities. This design allows researchers to collect data from multiple group members simultaneously and enables multiple trials per individual per session, greatly accelerating the rate at which robust cognitive datasets can be assembled in the wild.

A second advantage is that the apparatus only dispenses rewards when two conditions are met: the facial detection model must identify a study subject and the assigned touchscreen task must be completed. If an animal from a non-target species approaches and interacts with the apparatus, no reward is dispensed because the model does not detect a valid face. This feature ensures compliance with research permits and minimizes food loss to non-study organisms. For example, at our field site coatis (*Nasua narica*) occasionally approach the testing platforms, but under this design they cannot obtain food rewards and quickly lose interest. Although we cannot fully control which individuals choose to participate (see Limitations), we can actively exclude species and individuals from whom we do not wish to collect data.

A third key advantage is that the method is adaptable to multiple species and field sites. If a research group already has a large database of images or videos of study subjects, a site-specific facial recognition model can be trained immediately. Even without such a database, the more general model we trained using the Ultralytics YOLOv5 architecture can serve as a starting point. Because this model was trained with a looser concept of a capuchin face, it generalizes to other primate species and, in our tests, detected human, marmoset, and macaque faces. Used as we describe here, this general model can be deployed to assemble a video database that can then be used to train a species- and site-specific model, as we did for the second iteration of our facial recognition model (see **Section 3.2**). The physical testing apparatus is similarly exportable. It is relatively lightweight (about 35 lbs; 15.9 kg) and can be carried by a single person. All electronic components were selected for low cost and ease of replacement, and the two components with which subjects directly interact (the infrared touchscreen overlay and camera) were chosen for durability and affordability. The internal design is modular, so failure of one component does not require replacement of the entire system.

Taken together, these features position our method at the vanguard of efforts to understand cognition in ecologically relevant contexts while retaining a high degree of experimental control. They substantially reduce the gap between laboratory experimentation and field studies and move us closer to a long-sought goal: studying primate cognition in the wild with the rigor and precision typically reserved for the lab.

### 4.2 CapuchinAI: a versatile system for studying within- and between-individual variation in cognition in the wild

Touchscreen methods have emerged in recent years as highly versatile and robust tools for testing primate cognition (e.g., Martin et al., 2022). They allow researchers to collect standardized data on a wide array of cognitive traits within the same experimental framework, facilitating direct comparisons across tasks, individuals, and species. Bringing such methods to the field makes it possible to collect systematic data across various cognitive domains, rather than testing a single aspect of cognition in isolation, and greatly expands the range of experiments that can be implemented within a single study population. Traditionally, each new cognitive experiment has required habituation to a different apparatus. In contrast, the programmable nature of touchscreen experiments means that subjects need only habituate to a single smart testing station, which can then present a modular battery of lab-validated tasks targeting domains such as learning, memory, inhibitory control, flexibility, and social decision making.

Our goal is for CapuchinAI to serve as a flexible, long-term testing platform where tasks can be added, removed, or reordered in software without altering the hardware, and contingency rules can be used to scaffold individuals through tests of increasing difficulty at their own pace. The facial recognition software allows each subject to follow a personalized progression through the battery, automatically delivering the next task once predefined performance criteria are met. Because the system can run continuously, it supports longitudinal cognitive testing in the wild and lays the groundwork for comparative, population-level, and individual-level studies of cognitive variation and its ecological and fitness correlates.

This method can make a significant advance in the ability to measure inter-individual variation in cognition in the wild. Because cognition is context-dependent and developmentally malleable, deploying lab-based psychometric tools in natural settings will, for the first time, allow us to study cognitive variation in the fitness-relevant environments in which animal minds evolved (Horn et al., 2022). This, in turn, will help us address several central questions in primate cognition, including the heritability of cognitive traits (Hopkins et al., 2014), whether variation reflects a general intelligence “g” factor (Beran and Hopkins, 2018; Matzel and Sauce, 2017) or a more modular cognitive architecture (Amici et al., 2012), and which naturally varying environmental factors shape primate cognitive development. Because CapuchinAI is designed to track performance over time at the individual level, it allows us to view inter-individual variation not as a single snapshot, but as trajectories of cognitive change across days, months, and years.

### 4.3. Potential Limitations

The primary limitation of this approach is that it is easier to implement at some field sites and with some species than others. Our method is most readily adapted where primates are already habituated to specific areas used for cognitive testing. This pre-habituation, as with capuchins at Taboga and their familiarity with testing platforms, increases the likelihood that individuals will approach and interact with touchscreen boxes. The system also works best with species that show relatively low neophobia and high levels of self-motivated exploration, which again fits capuchins well. With an appropriate power supply, however, the apparatus can be left in place for extended periods without direct researcher supervision. Installing multiple units across the range of study groups or populations can promote passive habituation and allow individuals to engage with the system at their own pace.

A second limitation concerns participation and sampling. Because participation is entirely voluntary, we cannot guarantee a minimum number of trials per session or per individual, in contrast to many laboratory paradigms that use fixed trial numbers. We can set an upper limit on trials per individual, which helps prevent a small number of subjects from monopolizing rewards, but this does not solve the problem of low or uneven participation. In our pilot deployment, adult and subadult males were the most frequent users of the apparatus, although an adult female and a juvenile female also approached and investigated it before being displaced. Such patterns may introduce sampling biases. Running multiple stations simultaneously, in different parts of the home range, may help increase access and broaden participation across age–sex classes.

A final limitation relates to the use of food rewards. At Taboga, we provisioned capuchins with a food item they already encounter during natural foraging, provided quantities that should not alter foraging patterns (daily routes and time at specific locations have not changed in our longitudinal GPS data), and ensured that animals never observed researchers handling the rewards. Even with these precautions, some sites or species will not permit any level of provisioning. In those contexts, adapting our method would require tasks that are intrinsically rewarding or otherwise sufficiently engaging to motivate participation without food incentives.

## ACKNOWLEDGEMENTS

We thank our colleagues and collaborators for their insightful discussions that helped shape the development of this project. We are especially grateful to the members of the Social Cognition and Primate Behavior Lab at Emory University and the Capuchinos de Taboga project for their feedback and intellectual contributions. We would further like to thank the members of the Collective Cognition Lab at the University of Rochester, specifically Professor Dora Biro and graduate students Oviya Mohan and Rithwik Cherian, for their valuable suggestions and guidance. We also thank the editors and anonymous reviewers for their thoughtful comments and suggestions.

## ETHICS STATEMENT

This study was reviewed and approved by the relevant oversight committees, including the Institutional Animal Care and Use Committee (IACUC) at Emory University, and Costa Rica’s Sistema Nacional de Areas de Conservación (SINAC). Research was conducted according to protocols approved by IACUC, SINAC, Costa Rica’s National Technical University (UTN), granting agencies, and adhering to Costa Rican law. This research adhered to the American Society of Primatologists’ Principles for the Ethical Treatment of Non-human Primates.

